# Interpretable variational encoding of genotypes identifies comprehensive clonality and lineages in single cells geometrically

**DOI:** 10.1101/2024.07.04.602109

**Authors:** Hoi Man Chung, Yuanhua Huang

## Abstract

Clone assignment in single-cell genomics remains a challenge due to its diverse mutation macrostructures and many missing signals. Existing statistical methods, for the sake of numerical convergence, pose strong constraints on the form of predicted mutation patterns, so they easily identify sub-optimally fitted clones that overlook weak and rare mutations. To solve this problem, we developed SNPmanifold, a Python package that learns flexible mutation patterns using a shallow binomial variational autoencoder. The latent space of SNPmanifold can effectively represent and visualize complex mutations of SNPs (single-nucleotide polymorphisms) in the form of geometrical manifolds. Based on nuclear or mitochondrial SNPs, we demonstrated that SNPmanifold can effectively identify a large number of multiplexed donors of origin (k = 18) that all existing unsupervised methods fail and lineages of somatic clones with promising biological interpretation. Therefore, SNPmanifold can reveal insights into single-cell SNPs more comprehensively than other existing methods, especially in complex datasets.

## 1 Introduction

Single-cell genomics is the study of mutations in DNA sequences of individual cells using omics approaches. It has various applications [1, 2] in oncology, prenatal diagnosis, tissue mosaicism, immunology, organogenesis, embryogenesis, germline transmission, microbiology, and neurobiology, where single-cell genetic heterogeneity is prominent and informative either as markers [3] or pleiotropic mutations [4]. To investigate single-cell genetic heterogeneity, researchers developed various sequencing protocols [5, 6, 7, 8, 9] and associated statistical methods to analyze such data, particularly covering either or both of the two highly-coupled tasks: 1. Calling high-quality mutations, and 2. Assigning cell clones based on them.

For the first task of calling high-quality mutations, single-cell somatic variant callers encompass both nuclear-based and mitochondrial-based methods. Nuclear-based calling methods include Monovar [10] which detects variants with a high posterior probability of more than one allele in the cell population, and more recent methods by leveraging phasing with germline variants to reduce technical false positives [11, 12, 13]. Mitochondrial-based calling methods include mgatk by using variance-mean-ratio on raw allele frequency in the cell population [14], and MQuad by using BIC (Bayesian information criterion) to select variants supporting more than one clone [15]. For the second task of assigning cell clones, cells are grouped into genetic-based clusters, possibly considering the constraints of being compatible with phylogenetic trees. Example methods include Bayesian mixture models with Bernoulli noise (e.g. SCG Single-Cell Genotyper and BnpC) [16, 17], Cardelino that uses a putative phylogenetic tree as a prior and supports binomial noise for scRNA-seq data [18], and methods with higher computational efficiency for demultiplexing donors with germline variants [19, 20].

Nevertheless, almost all existing statistical methods are highly context-specific and pose strong constraints on the form of predicted mutation patterns for the sake of numerical convergence. Such strong constraints greatly reduce the flexibility of statistical models, so they easily underfit or overfit single-cell mutation patterns when hyperparameters, such as the number of clusters (k) of cells or the critical p-value for variant calling, are not well optimized. In most cases, these hyperparameters are not clear from experimental settings, thus researchers set them according to general model selection criteria such as BIC (Bayesian information criterion) or p-value<0.05. These general model selection strategies easily overlook informative mutations that are weak and rare, so we decided to develop a more reliable statistical method that can robustly identify weak and rare mutations which are consistent strong mutations normally revealed by other statistical methods.

Inspired by recent successes of VAE (variational autoencoder) [21], a kind of generative factor analysis neural network model, in solving numerous challenging biological problems, such as predicting cell transition [22, 23, 24] and integrating knowledge from different omics [25], we tried to apply VAE to identifying single-cell mutation patterns of SNPs. We realized that VAE with a well-optimized structure (upper right of Fig. 1) can informatively represent and visualize single-cell mutation patterns of SNPs as big or small, connected or disconnected geometrical manifolds. It can therefore serve as both a numerical and a visual guide for researchers to select sets of cells and co-mutations which align with mutation macrostructures, specifically mutation gaps and protruding lineages.

**Figure 1:**
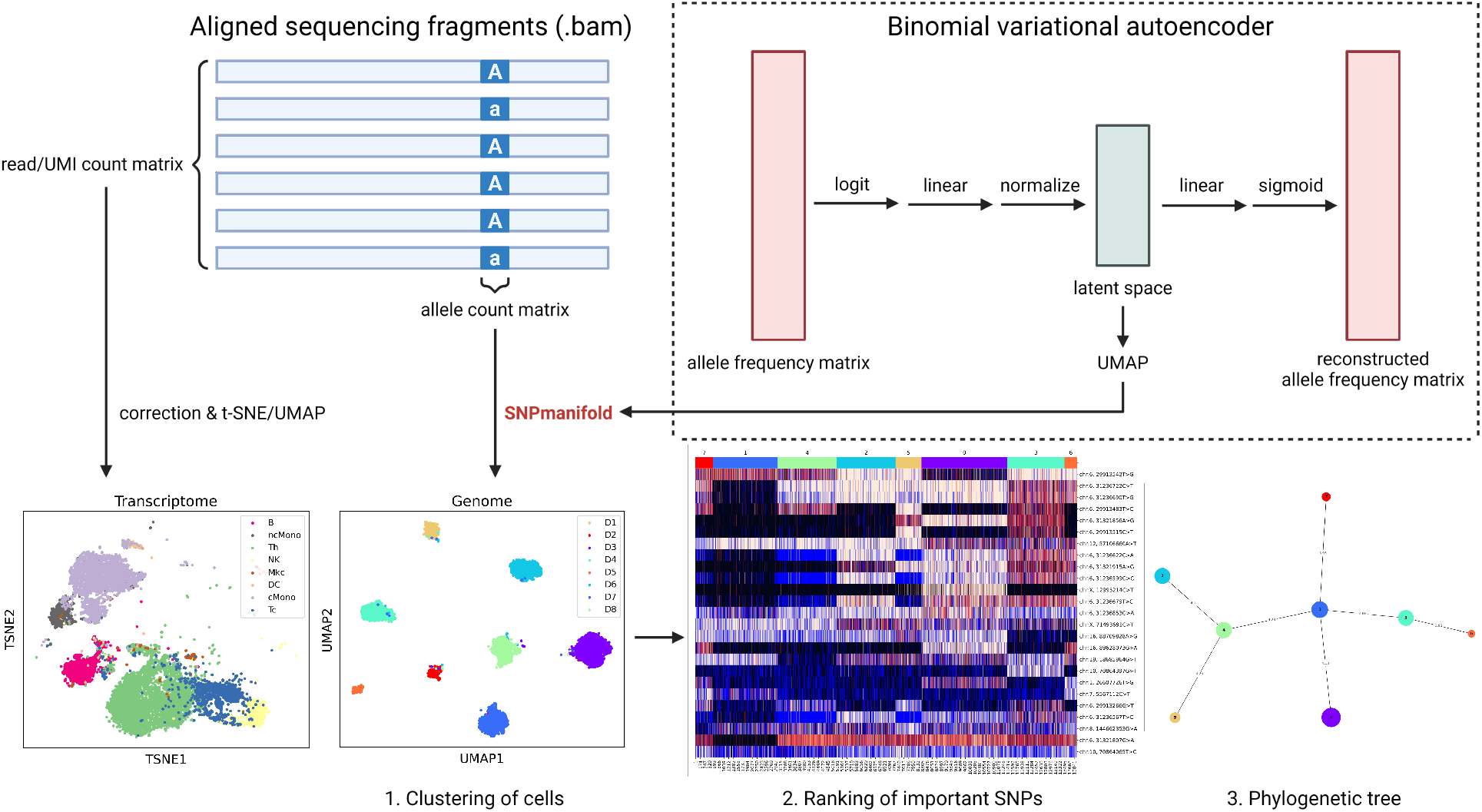
Schematic showing the workflow of SNPmanifold in a donor-multiplexing dataset (Donor8). After performing single-cell sequencing and obtaining a BAM file, researchers normally analyze the transcriptomic part by compiling a read/UMI count matrix and transforming it to an informative transcriptomic geometrical manifold using t-SNE or UMAP. SNPmanifold offers a genomic analog to this. After compiling an allele count matrix, SNPmanifold transforms it into an informative genomic geometrical manifold using a binomial variational autoencoder and UMAP. SNPmanifold then performs 3 downstream analyses on the genomic geometrical manifold: 1. Clustering of cells with similar genotypes (in latent space), 2. Ranking of important SNPs (via F-test), and 3. Constructing the phylogenetic tree (via agglomerative method). SNPmanifold can visualize biological insights for genomics in the same way t-SNE or UMAP can visualize for transcriptomics.

In this work, we introduce SNPmanifold (lower middle and right of Fig. 1), a Python package dedicated to convenient analysis and straightforward interpretation of the latent space in VAE (upper right of Fig. 1). SNPmanifold performs three downstream analyses on the genomic geometrical manifold: 1. Clustering of cells with similar genotypes (in latent space), 2. Ranking of important SNPs (via F-test), and 3. Constructing the phylogenetic tree (via agglomerative method). By learning a lower-dimensional (but not extremely low) embedding, these three standard tasks can benefit from effective strategies used in single-cell transcriptomic analysis, including exploration via visual examination and continuous space analysis such as graph-based clustering. Overall, SNPmanifold is a single-cell genetic analysis tool that adopts the philosophy of being clone-centric and exploration-friendly. In the following sections, we will demonstrate how SNPmanifold can reveal insights into single-cell clonality and lineages more comprehensively than other existing statistical methods, in diverse scenarios covering demultiplexing a large number of donors via germline variants and revealing somatic lineages via mitochondrial variants in cancer cell lines and primary human samples.

## 2 Results

### 2.1 Overview of SNPmanifold and the structure of variational autoencoder

VAE (variational autoencoder) is a kind of generative factor analysis neural network model that learns the generative process to reconstruct inputs from raw inputs. It typically consists of an encoder, a neural network with multiple progressively narrowing latent layers, and a decoder, a neural network with multiple progressively widening latent layers. Between the encoder and the decoder, there is one single narrowest hidden layer called the latent space, which theoretically captures the most informative features of raw input data. When the prior in the latent space is Gaussian, this model is called a deep latent Gaussian model or DLGM. Demonstrated by previous studies [22, 23, 24, 25] on VAE in solving other biological problems, this latent space can effectively identify macrostructures in the distribution of input data that are unclear in raw space and also correct missing values in individual input data, provided that the structure of VAE and its training strategy are well-optimized. This motivated us to adopt VAE with standard Gaussian prior as the backbone statistical model for SNPmanifold, a specialized strategy for detecting flexible single-cell mutation patterns and guiding model selection in single-cell mutation clustering.

To optimize the structure of VAE for single-cell genomics, we first replaced the Gaussian likelihood cost function in typical VAE with a binomial likelihood cost function which better describes the statistical distribution of allele counts in single-cell genomics data, particularly considering the coverage is generally low with a large proportion of zeros. We then examined VAEs with different numbers of hidden layers (LeakyReLU activation in between layers; Supplementary Fig. 2a), different values of *β* (strength of standard Gaussian prior; Supplementary Fig. 2b), and different values of D (numbers of latent dimensions; Supplementary Fig. 2c), on Vireo’s tutorial dataset (Donor4) [19], a scRNA-seq dataset which multiplexes 734 single cells with 1,905 SNPs from 4 donors (upper half of Fig. 2, Supplementary Fig. 2, 3). Upon investigation, we realized that shallow VAE with 1 hidden layer, *β* = 0, and D = 1/2 of the number of input SNPs, resolves single cells from 4 donors into 4 disconnected manifolds separated the best geometrically. In terms of visualizing the macrostructure of 4 donors (Fig. 2a, Supplementary Fig. 2a–c), this VAE visibly outperforms PPCA of raw allele frequency matrix, UMAP of raw allele frequency matrix, and other VAEs with larger numbers of hidden layers, larger values of *β*, and smaller values of D.

**Figure 2:**
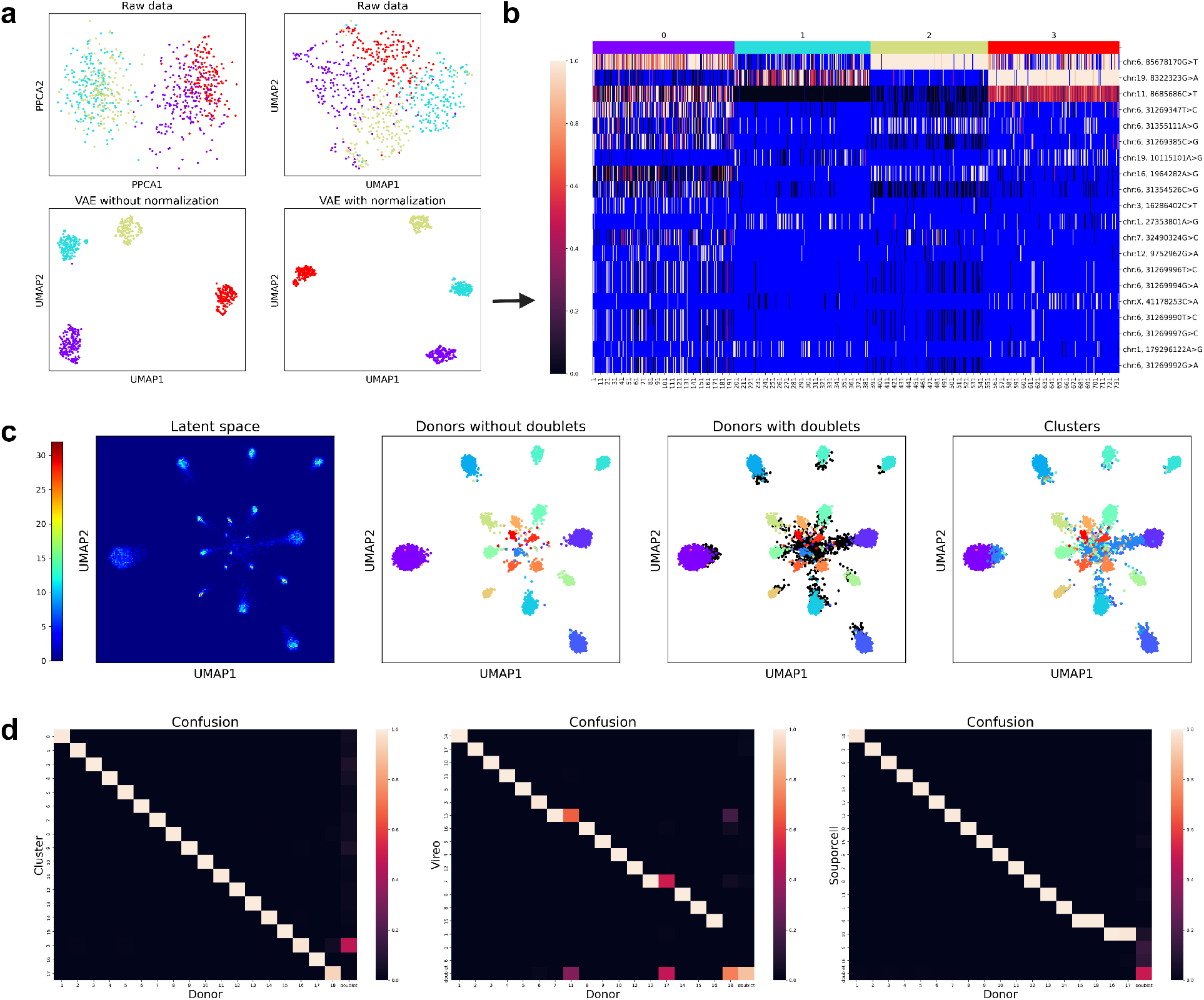
SNPmanifold demultiplexes scRNA-seq with a large number of donors accurately and visualizes them in disconnected manifolds. **a**, Genomic geometrical manifolds of Donor4 dataset in PPCA of the raw matrix, UMAP of the raw matrix, and UMAPs of the latent space in VAEs with 1 hidden layer with or without observed-SNP normalization (cell-specific division by the number of observed SNPs: sum(DP > 0)). VAE with 1 hidden layer and observed-SNP normalization geometrically separates 4 donors the best. **b**, Heatmap of allele frequency matrix of Donor4 dataset. The blue color indicates missing values (i.e. DP = 0). VAE resolves 4 donors into 4 clusters successfully despite many missing values. **c**, Density plot of the genomic manifold of Donor18 dataset, ground-truth donor labels with or without doublets, and SNPmanifold cluster labels on the manifold. There are 18 disconnected manifolds in the density plot corresponding to 18 donors, and Leiden clustering implemented in SNPmanifold identifies them successfully. **d**, Confusion matrices between donor labels and our cluster labels, donor labels and Vireo labels, and donor labels and Souporcell labels. The values of each column sum to 1. The accuracy between donor labels and cluster labels reaches 99.3% while Vireo and Souporcell fail to identify donors accurately.

To account for random missing signals due to technical artifacts across single cells, we further introduced observed-SNP normalization (cell-specific division by the number of observed SNPs: sum(DP > 0)) between the linear transformation in the encoder and the latent space. It visibly increases manifold separation for 4 donors compared to that without observed-SNP normalization (Fig. 2a, Supplementary Fig. 2d). Up to this point, we thought the structure was well-optimized for single-cell genomics, then we adopted VAE with this structure (upper right of Fig. 1) for SNPmanifold and the rest of the results section. Of note, while the model structure was mainly assessed via visual examination of the manifolds, the same conclusion is supported by the quantitative accuracy of the clustering results in the next paragraph. All three downstream analyses of SNPmanifold — 1. Clustering of cells with similar genotypes (in latent space), 2. Ranking of important SNPs (via F-test), and 3. Constructing the phylogenetic tree (via agglomerative method) — starts with the latent space in this VAE.

### 2.2 SNPmanifold projects multiplexed donors of origin to multiple disconnected manifolds

After finalizing the structure of VAE in SNPmanifold, we first quantitatively assessed its performance in the above Donor4 dataset for donor assignment. K-means clustering of the latent space (Supplementary Fig. 3a) identifies 4 donors successfully with good clustering metrics (distortion and silhouette score) and an accuracy of 99.8% compared to donor labels assigned by Vireo [19] without reference to genotypes, a published donor-demultiplexing statistical method (Supplementary Fig. 3b).

Next, we applied SNPmanifold to Demuxlet’s benchmarking dataset (Donor8) [26], a scRNA-seq dataset that multiplexes 13,939 PBMCs (peripheral blood mononuclear cells) with 929 SNPs from 8 donors (lower middle and right of Fig. 1, Supplementary Fig. 1). VAE resolves single cells from 8 donors into 8 disconnected manifolds successfully (lower middle of Fig. 1), and k-means clustering of the latent space (lower middle of Fig. 1, Supplementary Fig. 1a) identifies them successfully with good clustering metrics (distortion and silhouette score) and an accuracy of 99.7% compared to donor labels assigned by Demuxlet [26] with reference to genotypes, a published donor-demultiplexing statistical method (Supplementary Fig. 1b).

Lastly, we applied SNPmanifold to a challenging dataset (Donor18) [27], a scRNA-seq dataset which multiplexes 9,436 iPSCs (induced pluripotent stem cells) with 864 SNPs from 18 donors (Fig. 2c, d, Supplementary Fig. 4, 5). Despite severely imbalanced sample sizes across donors (67 to 1,519 cells; Supplementary Fig. 4c), VAE resolves single cells from 18 donors into 18 disconnected manifolds successfully (Fig. 2c), and Leiden clustering of the latent space (Fig. 2c, Supplementary Fig. 4a) identifies them successfully with an accuracy of 99.3% compared to donor labels assigned by Vireo [19] with reference to genotypes, a published donor-demultiplexing statistical method (Fig. 2d). SNPmanifold outperforms Vireo [19] without reference to genotypes and Souporcell [20] without reference to genotypes, two published donor-demultiplexing statistical methods, where both Vireo and Souporcell erroneously grouped two donors into one cluster but SNPmanifold placed each of the 18 donors into a separate cluster (Fig. 2d, Supplementary Fig. 5). From the viewpoint of model selection in single-cell mutation clustering, the latent space of our VAE enables direct visual examination of the model via the consistency between its cluster label and disconnected manifolds. Hence, our VAE can predict the success of Vireo with reference to genotypes (Fig. 2C, second panel) and the failures of Vireo and Souporcell without reference to genotypes (Fig. 2D) in this challenging dataset, because the first clustering result (Fig. 2c) align perfectly with disconnected manifolds of the latent space while the latter two clustering results (Supplementary Fig. 5) do not.

### 2.3 SNPmanifold projects lineages of somatic clones to multiple disconnected tree-like manifolds

After initial successes of SNPmanifold in 3 donor-multiplexing datasets, we would like to demonstrate the capability of SNPmanifold in identifying lineages of somatic clones within individual cell lines. For convenience in benchmarking our method, we focused only on mitochondrial SNPs because they have fewer positions, higher sequencing depths, and higher mutation rates. Also, we could avoid the use of different SNP-calling methods for the nuclear genome which may confound the results. Briefly, for each dataset, SNPmanifold performs cell and variant pre-filtering on raw input matrices (removing lowly covered variants and cells, and selecting highly variable variants), and then inputs filtered SNPs into the VAE model to obtain a latent embedding, where it then performs clustering, detects clonal variants via F-tests between cell clusters, and reconstructs a phylogenetic tree in an agglomerative way (see more details in Methods).

We first applied SNPmanifold to mtscATAC-seq’s benchmarking dataset (TF1 GM11906) [14], a singlecell mitochondrial ATAC-seq dataset which mixes 1001 single cells with 56 mitochondrial SNPs from two hematopoietic cell lines (TF1 and GM11906 cell lines; Fig. 3a–c, Supplementary Fig. 6). VAE resolves single cells into 8 disconnected manifolds (Fig. 3b). If we consider each disconnected manifold to be one separate lineage, SNPmanifold identifies a total of 8 lineages: 4 correspond to TF1 cell line (Fig. 3a and 3b, bottom right), 3 correspond to GM11906 cell line (Fig. 3a and 3b, bottom left), and 1 corresponds to doublets of the two cell lines (cluster 22; Fig. 3a), according to a set of 47 nearly homozygous SNPs (16126T>C, 150T>C, 16298T>C, etc.; Fig. 3a, Supplementary Fig. 6a). Out of the 4 lineages from TF1 cell line, 1 corresponds to wild type and 1 corresponds to 309C>T. The remaining 2 share the same mutation 7789G>C&8002C>T; 1 lineage possesses additional mutation 3901G>A, while the other lineage is tree-like and possesses either no additional mutation or one additional mutation 13708G>A or 4037G>A, in different parts of this lineage. On the other hand, for the 3 lineages from the GM11906 cell line, 1 corresponds to wild type, 1 corresponds to 8344A>G, and 1 corresponds to 8202T>C&8344A>G. All the lineages identified by SNPmanifold are consistent with the major variants reported by the original paper which used their own statistical method, mgatk [14], to select important SNPs. In terms of phylogeny, SNPmanifold constructs a cluster phylogenetic tree (Fig. 3c) in the latent space which shows a distinct separation between TF1 cell line clusters on top and GM11906 cell line clusters at bottom, connected by the cluster of doublets in the middle. Altogether, SNPmanifold identifies mutation patterns of two orders in this dataset, one big germline mutation between the two cell lines and many small somatic mutations within each cell line.

Next, we applied SNPmanifold to MQuad’s benchmarking dataset (MKN45) [28, 15], a scDNA-seq dataset which contains 5,199 single cells with 322 SNPs from MKN45 human gastric cancer cell line (Fig. 3d–f, Supplementary Fig. 7). VAE resolves single cells into 2 disconnected manifolds, 1 upper-right small manifold and 1 lower-left large tree-like manifold (Fig. 3e). The upper-right small manifold represents a simple lineage with mutation 2393C>T&8368G>A, while the lower-left large manifold is tree-like and represents a complex lineage with a varying major haplotype of 6 SNPs (1841T>C, 14552G>A, etc.) as well as occasional additional mutation 3436G>A, 8940C>T, 7236G>A, etc. (Fig. 3d, e, Supplementary Fig. 7a). Projecting clone labels assigned by MQuad [15], a published mitochondrial clone assignment method, or another simple way using a set of 3 occasional mutations onto the latent space (Fig. 3e, Supplementary Fig. 7c), we realized that they actually correspond to different ways of partitioning the same latent space. From the viewpoint of model selection in single-cell mutation clustering, both clone assignment results are reasonable because they both highlight the major separation between the upper-right small manifold, which has a distinct copy number variation as reported in MQuad paper [15], and the lower-left large tree-like manifold. However, both clone assignment results are underfitted because MQuad ignores occasional mutations with low BICs and the set of 3 occasional mutations ignores the major haplotype MQuad highlights. SNPmanifold, in contrast, captures full conditional dependence between the major haplotype and different occasional mutations, and represents it as one continuous tree-like manifold which describes the gradual, partially-synchronized evolution of different SNPs. SNPmanifold then constructs a cluster phylogenetic tree (Fig. 3f) in the latent space to summarize these complex evolutionary dynamics for easy and straightforward human interpretation.

**Figure 3:**
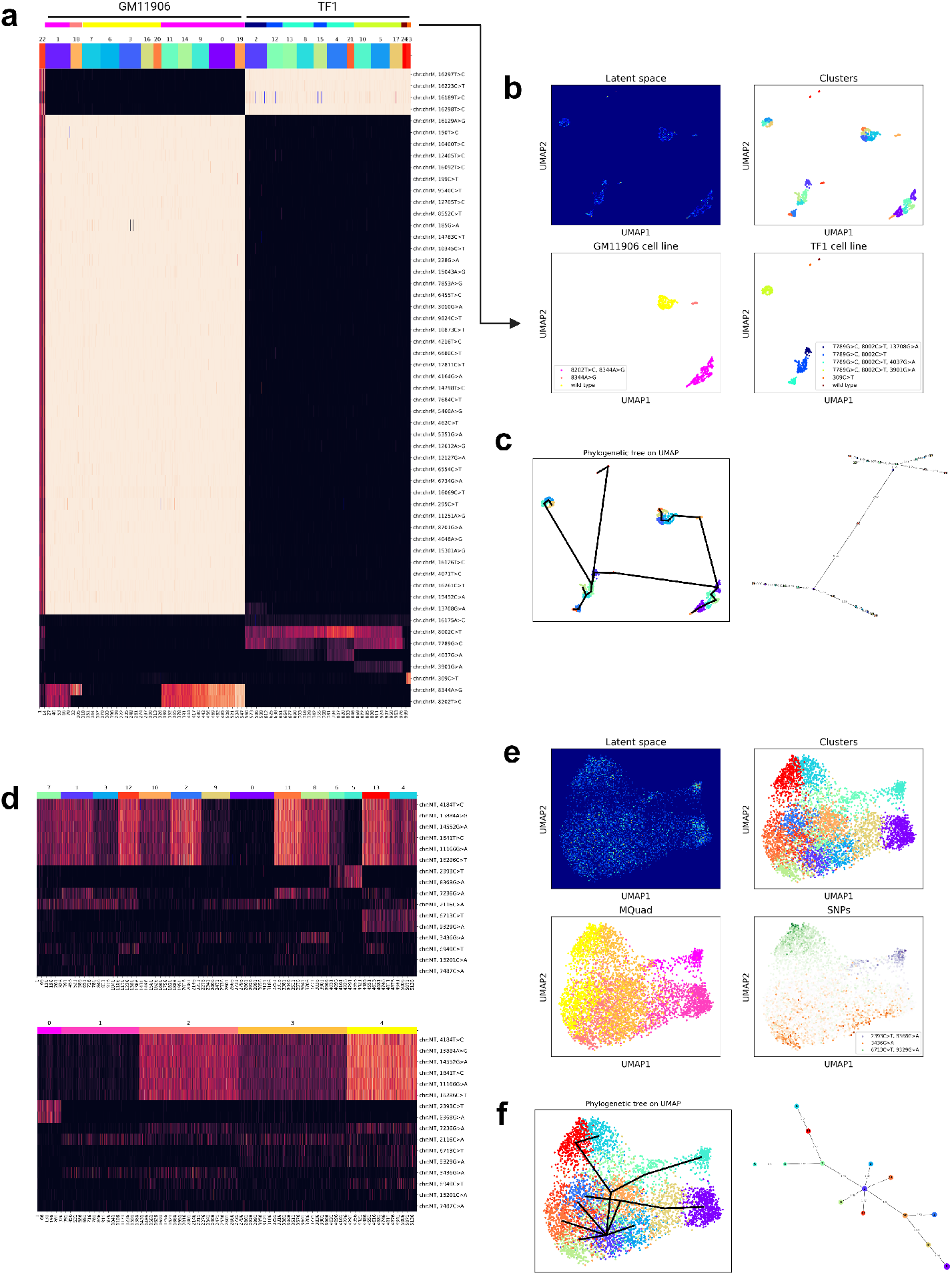
SNPmanifold identifies interpretable lineages and the phylogeny within cell lines by using mitochondrial variants. **a**, Heatmap of allele frequency matrix of TF1 GM11906 dataset. VAE highlights 2 major genotypes corresponding to 2 cell lines, and additional somatic mutations within each cell line. **b**, Density plot of the genomic manifold of TF1 GM11906 dataset, cluster labels on the manifold, coarse genotypes within each cell line on the manifold. Some genotypes are connected together in one manifold (one connected lineage), while others are separated in different manifolds (separate lineages). **c**, Phylogenetic tree of the manifold of TF1 GM11906 dataset. The tree constructed by SNPmanifold clearly separates 2 cell lines with doublets (cluster 22) placed in the middle. **d**, Heatmaps of allele frequency matrix of MKN45 dataset, with cells sorted by SNPmanifold (upper) or MQuad (lower). SNPmanifold assigns cell clones with more conditionally-dependent occasional mutations compared to MQuad. **e**, Density plot of the genomic manifold of MKN45 dataset, cluster labels and MQuad labels on the manifold, allele frequency of 3 occasional mutations on the manifold. MQuad labels and the 3 occasional mutations actually correspond to different ways of partitioning the same manifold. This means that the genomic manifold is consistent with both clone assignment results while being more comprehensive than either of them. **f**, Phylogenetic tree of the manifold of MKN45 dataset.

### 2.4 SNPmanifold reveals correlations between cellular phenotypes and phylogeny of genotypes

Encouraged by the successes of SNPmanifold in identifying lineages of somatic clones within individual cell lines, we would like to further demonstrate its capability in identifying lineages of somatic clones within primary human samples. Again, for convenience in benchmarking our method, we focused only on mitochondrial SNPs because they have fewer positions, higher sequencing depths, and higher mutation rates. Also, we could avoid the use of different SNP-calling methods for the nuclear genome which may confound the results. Briefly, for each dataset, SNPmanifold performs cell and variant pre-filtering on raw input matrices (removing lowly covered variants and cells, and selecting highly variable variants), and then inputs filtered SNPs into the VAE model to obtain a latent embedding, where it then performs clustering, detects clonal variants via F-tests between cell clusters, and reconstructs a phylogenetic tree in an agglomerative way (see more details in Methods).

We first applied SNPmanifold to MAESTER’s benchmarking dataset (BPDCN) [9], a single-cell mitochondrial RNA-seq dataset that contains 9,204 PBMCs (peripheral blood mononuclear cells) with 274 SNPs from a patient with clonal hematopoiesis (Fig. 4a–e, Supplementary Fig. 8, 9). VAE resolves single cells into many disconnected manifolds (Fig. 4b): 6 of them represent lineages with missing transcripts ND1, ND2, ND5, CYTB, ND1&ND2, ND1&ND5, and 3 of them represent lineages with mutations 2593G>A, 683G>A, 6205G>A&9164T>C, respectively (Fig. 4a, b, Supplementary Fig. 8a). In terms of phylogeny, the cluster phylogenetic tree constructed by SNPmanifold suggests the existence of 3 major common ancestors (clusters 4, 12, 19), each of which directly evolves into at least 5 other clusters. Projecting cell-type labels onto the latent space or the phylogenetic tree (Fig. 4c, d, Supplementary Fig. 8c, 9a) straightforwardly reveals correlations between cell types and phylogeny of genotypes: most HSPCs (hematopoietic stem and progenitor cells) and myeloid descendants localize around the common ancestor (cluster 19) on top, most B cells localize around the common ancestor (cluster 4) in the middle, while T cells and NK cells are prevalent around all 3 common ancestors. Projecting TRB-clonotype labels onto the latent space or the phylogenetic tree (Fig. 4c, e, Supplementary Fig. 8d, 9b) further reveals strong correlations between certain TRB-clonotype and certain clusters, such as CASSFRQGYNEQFF in cluster 29 with mutation 683G>A and CASSLEWGNPSTYEQYF in cluster 31 with mutation 6205G>A&9164T>C.

**Figure 4:**
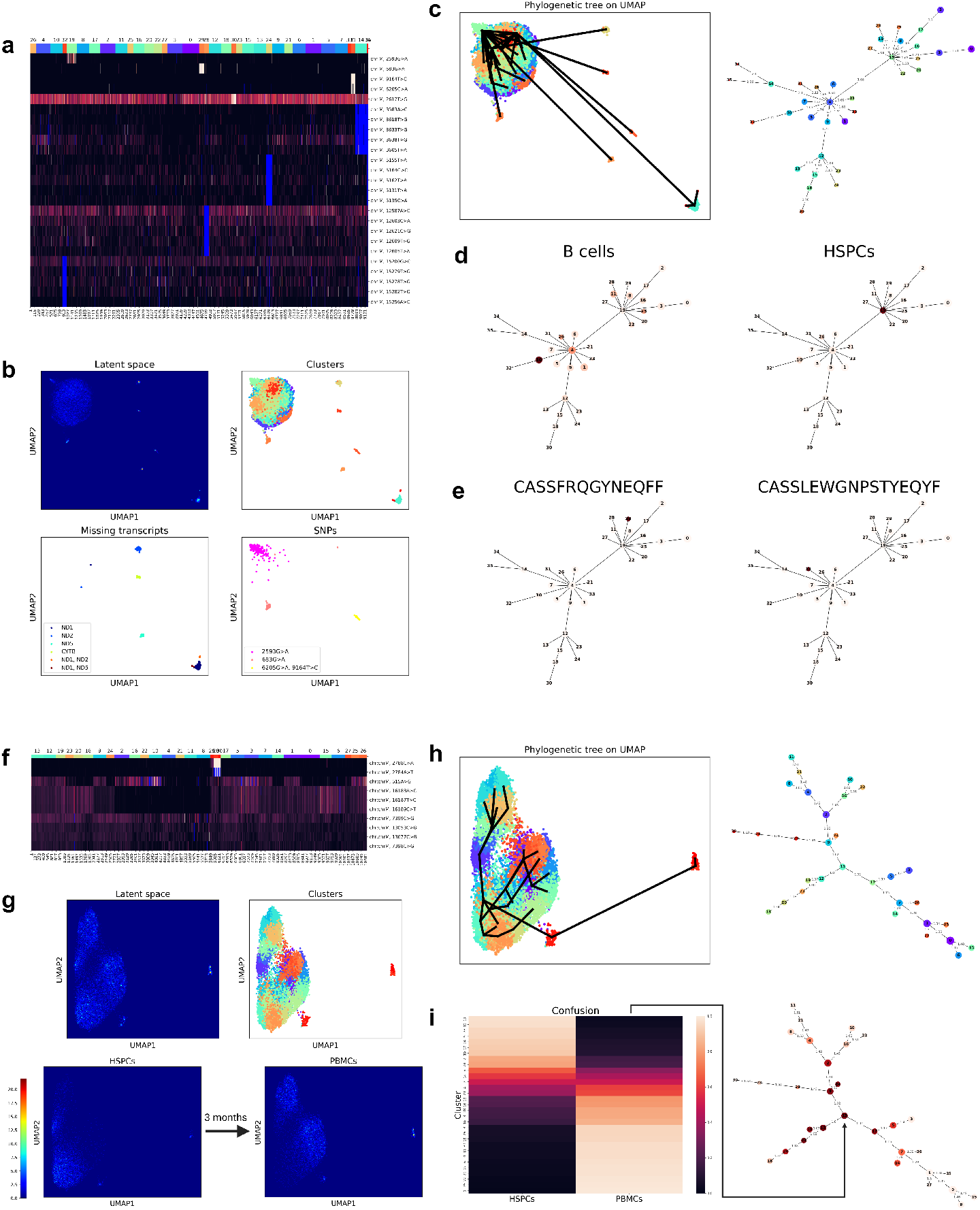
SNPmanifold reveals biological insights of somatic clones in primary human samples. **a**, Heatmap of allele frequency matrix of BPDCN dataset. Blue color indicates missing values (i.e. DP = 0). VAE highlights some somatic mutations and some missing transcripts. **b**, Density plot of genomic manifold of BPDCN dataset, cluster labels on the manifold, coarse genotypes and missing transcripts on the manifold. **c**, Phylogenetic tree of the manifold of BPDCN dataset. **d**, Relative abundance of 2 cell types on phylogenetic tree of the manifold of BPDCN dataset. Deeper color indicates higher abundance. **e**, Relative abundance of 2 TRB clonotypes on phylogenetic tree of the manifold of BPDCN dataset. Deeper color indicates higher abundance. **f**, Heatmap of allele frequency matrix of HSPC PBMC dataset. Blue color indicates missing values (i.e. DP = 0). VAE highlights several somatic mutations. **g**, Density plot of genomic manifold of HSPC PBMC dataset, cluster labels on the manifold, HSPCs/PBMCs on the manifold. The manifold captures the temporal evolution of genotypes from HSPCs to PBMCs after 3 months. **h**, Phylogenetic tree of the manifold of HSPC PBMC dataset. **i**, Confusion matrix between HSPCs/PBMCs and cluster labels of HSPC PBMC dataset, the relative abundance of cell types on phylogenetic tree of the manifold. Deeper color indicates a higher abundance of HSPCs and a lower abundance of PBMCs. There is a clear gradual color change from deep colors (HSPCs) in the middle of the tree (cluster 13) to bright colors (PBMCs) on the periphery of the tree. (Note: Larger figures can be found in the code availability section.)

Lastly, we applied SNPmanifold to mtscATAC-seq’s another benchmarking dataset (HSPC PBMC) [14], a single-cell mitochondrial ATAC-seq dataset which contains 4,339 HSPCs (hematopoietic stem and progenitor cells) and 6,597 PBMCs (peripheral blood mononuclear cells) with 100 SNPs from a healthy donor (Fig. 4f–i, Supplementary Fig. 10). PBMCs are collected 3 months after HSPCs. VAE resolves single cells into 4 disconnected manifolds (Fig. 4g): 1 corresponds to wild type, 1 corresponds to weak mutation 16187 16189TAC>CCT, 1 corresponds to weak mutation 2788C>A, and 1 corresponds to strong mutation 2784A>T&2788C>A (Fig. 4f, g, Supplementary Fig. 8a). Projecting HSPCs/PBMCs labels onto the latent space or the phylogenetic tree (Fig. 4g–i) straightforwardly reveals correlations between cell types and phylogeny of genotypes from HSPCs to PBMCs after 3 months. Firstly, from the latent space (Fig. 4g), 4 disconnected manifolds of HSPCs evolve into 4 disconnected manifolds of PBMCs after 3 months, suggesting an inheritance of characteristic ancestral mutations. Secondly, the genetic difference between HSPCs and PBMCs arises primarily from a systematic decrease in weak mutation 13053C>G&13677C>G (Fig. 4f, Supplementary Fig. 10a). Thirdly, from the phylogenetic tree (Fig. 4i), HSPCs mostly localize in the middle while PBMCs mostly localize on the periphery, in line with the biological theory where HSPCs with similar genotypes divide into PBMCs with more dissimilar genotypes. Fourthly, wild-type HSPCs (cluster 2) divide into 2 sub-lineages of PBMCs (clusters 4, 16), while HSPCs with weak mutation 16187 16189TAC>CCT (cluster 13) divide into 4 sub-lineages of PBMCs (clusters 1, 3, 18, 26).

## 3 Discussion

The primary motivation of our work was to develop a flexible statistical model that can identify comprehensive genetic clonality and lineages in single cells without posing strong constraints on the form of predicted mutation patterns. Before our SNPmanifold, almost all existing statistical methods in single-cell genomics fall into one of the two main categories: 1. Donor-demultiplexing methods which identify a small fixed number of origin haplotypes, or 2. One-by-one SNP-selection methods which assume different variants to be largely independent. Both types of methods pose strong constraints on the form of predicted mutation patterns so they easily underfit or overfit when their underlying assumptions are violated. This problem of sub-optimal fitting is further exacerbated when their hyperparameters, such as the number of clusters (k) or the critical p-value, are not clear from experimental settings so researchers have to guess with arbitrarity using general model selection rules. To solve this problem, we decided to try binomial VAE (variational autoencoder), a statistical model which can detect flexible mutation patterns yet being numerically robust.

For the architecture of the VAE framework, a recent paper [29] also working on VAE for single-cell genomics suggested that the optimal structure of VAE should be 5 hidden layers with LeakyReLU activation. After a brief trial of their VAE on the Donor4 dataset (rightmost of Supplementary Fig. 2a), we realized our VAE provides better visualization of the latent space while keeping high clustering accuracy. This can be due to some of the unconventional modifications we made, such as removing LeakyReLU activation, reducing the number of hidden layers to 1, and adding observed-SNP normalization (cell-specific division by the number of observed SNPs: sum(DP > 0)).

Conceptually, we view our VAE to be a constrained form of ordinary factor analysis with the following modifications: 1. Linear encoder, 2. Perturbation to the latent space in training, and 3. Observed-SNP normalization. Firstly, unlike factor analysis which optimizes cell-specific latent factors independently, our VAE optimizes a linear encoder that performs amortized inference on cell-specific latent factors using the same linear function for all cells. This confers two important advantages: 1. Cells with similar genotypes must be mapped to neighbors in the latent space as a connected manifold, and 2. Overfitting of cell-specific latent factors to outlier cells is prevented by having the same linear function for all cells. Secondly, perturbation to the latent space in training helps escape unstable local optima and converge to a desirable stable global optimum, which enjoys the merits of being an approximate Bayesian method. Thirdly, observed-SNP normalization (cell-specific division by the number of observed SNPs: sum(DP > 0)) reduces dispersion of the same genotype in the latent space by explicitly correcting random missing signals due to technical artifacts across single cells. Altogether, these 3 modifications help our VAE to stably learn non-overfitted geometrical manifolds which can visualize single-cell mutation macrostructure clearly.

In the results section, our VAE has demonstrated its capability in visualizing mutation patterns of different orders, including weak mutations with low allele frequency, strong mutations with high allele frequency, missing transcripts, and haplotypes, altogether coherently and integratively in a 2D UMAP. By projecting the allele frequency of SNPs onto the 2D UMAP (upper right of Fig. 3e, Supplementary Fig. 6a, 7a, 8a, 10a), conditional dependence between mutations of different SNPs becomes readily visible to researchers. This is an unprecedented achievement, because existing statistical methods in single-cell genomics focus only on either big haplotypes or mutations of independent SNPs, ignoring possible interactions between the two. Compared to them, SNPmanifold presents a more complete picture of the underlying single-cell genetic heterogeneity in the dataset. With the manifestation of mutation gaps and protruding lineages in 2D UMAP, researchers can rapidly identify an appropriate number of clusters (k) which are otherwise ambiguous, or disconnected lineages and rare clones which are overlooked by other methods. SNPmanifold can also be used for the diagnosis of sub-optimal clustering (e.g. donor assignment) and provide guidance to post hoc finetuning of clustering results which does not align perfectly with mutation gaps due to convergence problems or slight violation of model assumptions.

In terms of single-cell phylogenetics, we adopted a strategy to focus on clone level instead of each individual cell. We think the cluster phylogenetic tree in our SNPmanifold extends more naturally to a high number of cells than conventional bifurcating phylogenetic tree, for two reasons. Firstly, our cluster phylogenetic tree can represent phylogenetic multifurcation which is common when the number of cells is high, but conventional bifurcating phylogenetic tree cannot. Secondly, the conventional bifurcating phylogenetic tree has poor analyzability and human-interpretability when it scales linearly to a high number of cells, but our cluster phylogenetic tree scales linearly to the number of clusters (k) which is much smaller and often determinable from 2D UMAP of the latent space. In practice, we think cluster phylogenetic tree and 2D UMAP are good complements to each other and should be interpreted together (Fig. 3c, f, 4c, h). For 2D UMAP, the cluster phylogenetic tree, which is constructed in the full-dimensional latent space, can compensate for inherent tradeoffs in 2D UMAP visualization such as loss of high-dimensional features and unguaranteed relative positions between disconnected manifolds. For cluster phylogenetic tree, 2D UMAP can offer information about big mutation gaps, therefore researchers can perform post hoc pruning on the cluster phylogenetic tree to obtain multiple small cluster phylogenetic trees representing disconnected lineages.

At last, we would like to outline some potential usage of our Python package, SNPmanifold, in future biomedical research. After obtaining single-cell sequencing results, users can use their own preferred method, such as cellsnp-lite [30], to compile allele count matrices (AD matrix, DP matrix) and input them to SNPmanifold. After training, SNPmanifold will output the latent space and its downstream analyses including clustering results of cells, 2D UMAP, heatmap of allele frequency matrix, cluster phylogenetic tree, etc. In short, SNPmanifold offers an integrated pipeline for analyzing and visualizing general single-cell genetic heterogeneity. Compared to existing statistical methods, SNPmanifold is more sensitive to weak and rare mutations so it can reveal additional biological insights in the same experimental sequencing data based on them. SNPmanifold can be particularly useful in donor demultiplexing and inferring tumour evolution, where rare mutations prevail due to imbalance clone size, or mitochondrial lineage tracing, where weak mutations are very informative.

## 4 Methods

### 4.1 Binomial variational autoencoder

VAE (variational autoencoder) is a kind of generative factor analysis neural network model that learns the generative process to reconstruct inputs from raw inputs. When the prior in the latent space is Gaussian, this model is called a deep latent Gaussian model or DLGM. SNPmanifold uses a shallow binomial VAE (variational autoencoder) with standard Gaussian prior as the backbone statistical model to identify singlecell mutation macrostructure from sequencing results. It takes a cell by SNP allele frequency matrix (AF matrix = AD*/*DP: count of the alternative allele divided by the total count of all alleles) as input. The structure of our VAE (upper right of Fig. 1) consists of logit transformation followed by linear transformation and observed-SNP normalization (cell-specific division by the number of observed SNPs: sum(DP > 0)) in encoder with parameters *ϕ*, and linear transformation followed by sigmoid transformation in decoder with parameters *θ*. After all transformations, it outputs a reconstructed allele frequency matrix, as follows

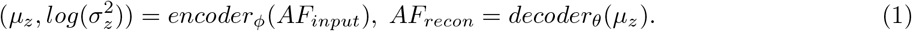

The cost function to optimize is the weighted sum (weight = *β*) of the negative log of binomial likelihood and the KL term in ordinary VAE:

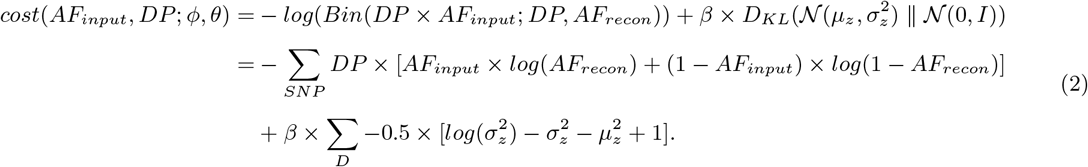

We optimized hyperparameters using the Donor4 dataset (Supplementary Fig. 2b, c) and set *β* = 0 and z = 1*/*2 of the number of input SNPs by default. We then train VAE for 2000 epochs by default using Adam optimizer (learning rate: 0.0001 and weight decay: 0) and encode the input allele frequency matrix into the latent space using the trained encoder. After that, SNPmanifold performs 3 downstream analyses on the genomic geometrical manifold in the latent space of VAE: 1. Clustering of cells with similar genotypes, 2. Ranking of important SNPs, and 3. Constructing the phylogenetic tree (lower middle and right of Fig. 1).

### 4.2 Pre-filtering of SNPs and cells

Before extracting the latent space using VAE, SNPmanifold first performs three pre-filtering steps on input SNPs and cells: 1. Number of observed SNPs for each cell, 2. Mean coverage of each SNP, and 3. Logit-variance of each SNP. SNPmanifold will provide plots on these three quantities to aid users in choosing cut-off for pre-filtering of high-quality SNPs and cells, based on elbow points in plots and overall sequencing qualities.

### 4.3 Clustering of cells with similar genotypes in the latent space

For clustering of cells with similar genotypes, SNPmanifold supports k-means clustering and Leiden clustering (graph-based clustering in SCANPY [31]) in either full-dimensional latent space or 3D UMAP [32]. The default method is Leiden clustering in 3D UMAP. Users should choose the clustering result that aligns better with mutation macrostructures, specifically mutation gaps and protruding lineages, visualized in 2D UMAP [32].

### 4.4 Ranking of important SNPs according to cell clusters

For ranking of important SNPs, SNPmanifold uses the inferred cell clonal clusters as input and prioritizes SNPs via an ANOVA test for each SNP. Specifically, it first computes F-tests between each cluster and the bulk (i.e. all cells) for each SNP and then ranks SNPs from the lowest p-value to the highest p-value (Supplementary Fig. 1c, 3c, 4b, 6b, 7b, 8b, 10b). Maximum F-statistic is capped at 20 and minimum p-value is capped at 10^−16^ for computational reasons. In the heatmap of the allele frequency matrix, SNPs are shown in the order of increasing p-value for visualization of important SNPs on top.

### 4.5 Constructing the phylogenetic tree in the full-dimensional latent space

For constructing the phylogenetic tree, SNPmanifold leverages an agglomerative strategy to generate a lineage graph. Specifically, it first computes by default the shortest 100 pairwise distances between cells from each pair of clusters in the full-dimensional latent space, and then iteratively connects the pair of clusters with the shortest average pairwise distance. In the end, all clusters will be connected into one acyclic graph which is the cluster phylogenetic tree with maximal parsimony in the full-dimensional latent space. In default visualization, the size of each node represents the number of cells in each cluster, and the number on each edge represents the distance between two clusters in the latent space.

### 4.6 Data availability

All datasets used in this study are previously published datasets. Donor4 [19] is downloaded from Vireo’s GitHub repository as SNP-count matrices. Donor8 [26] is downloaded from GEO sample GSM2560248 as FASTQs. Donor18 [27] is downloaded from ENA biosample SAMEA6833385 as FASTQs. TF1 GM11906 [14] is downloaded from GEO sample GSM4238432 as FASTQs. MKN45 [28] is downloaded from 10x Genomics as BAM. BPDCN [9] is downloaded from GEO sample GSM5534706 as SNP-count matrices. HSPC PBMC [14] is downloaded from GEO sample GSM4472965 and GSM4472967 as FASTQs. For the aforementioned FASTQs, we first aligned them to hg19 or hg38 to BAM and then compiled SNP-count matrices using cellsnp-lite [30]. Processed input matrices for SNPmanifold can be downloaded from https://github.com/StatBiomed/SNPmanifold/tree/main/data.

## 5 Code availability

SNPmanifold is a standard and open-source Python package with source code freely available at https://github.com/StatBiomed/SNPmanifold. For reproducibility, the analysis notebooks and figures are also available in this repository.

