## supplementary figures for "Interpretable variational encoding of genotypes identifies comprehensive clonality and lineages in single cells geometrically"

Supplementary 1

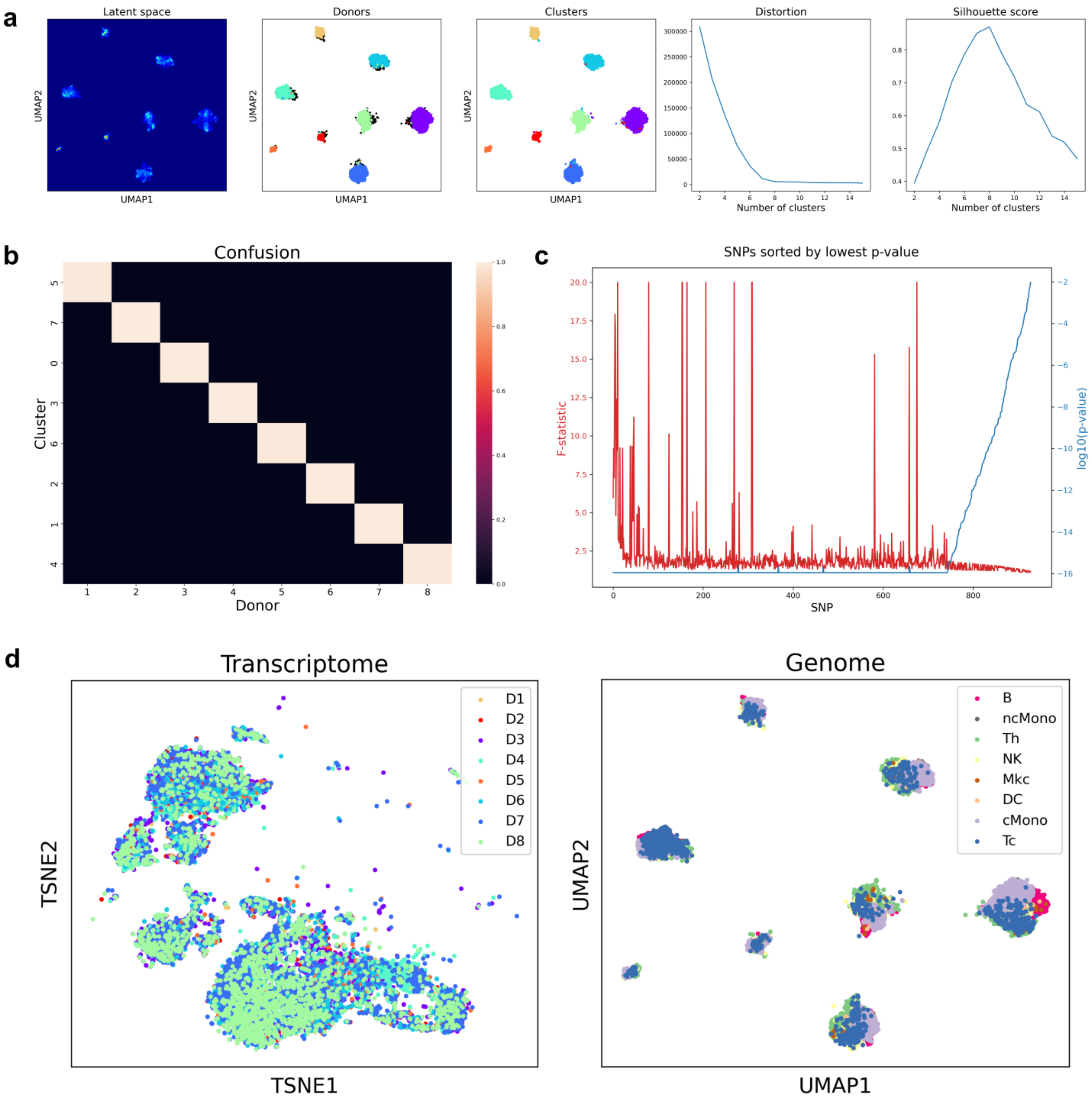


**a,** Density plot of genomic manifold of Donor8 dataset, donor labels and cluster labels on the manifold, distortion and silhouette scores of k-means clustering. There are 8 disconnected manifolds in the density plot corresponding to 8 donors, and k-means clustering identifies them successfully with good clustering metrics. **b,** Confusion matrix between donor labels and cluster labels of Donor8 dataset. The accuracy reaches 99.7%. **c,** F-statistics and p-values of the SNPs of Donor8 dataset ranked by SNPmanifold. **d,** Cluster labels on transcriptomic t-SNE and cell-type labels on genomic manifold. These two manifolds are largely independent of each other, meaning that they can offer biological insights from different perspectives.

Supplementary 2

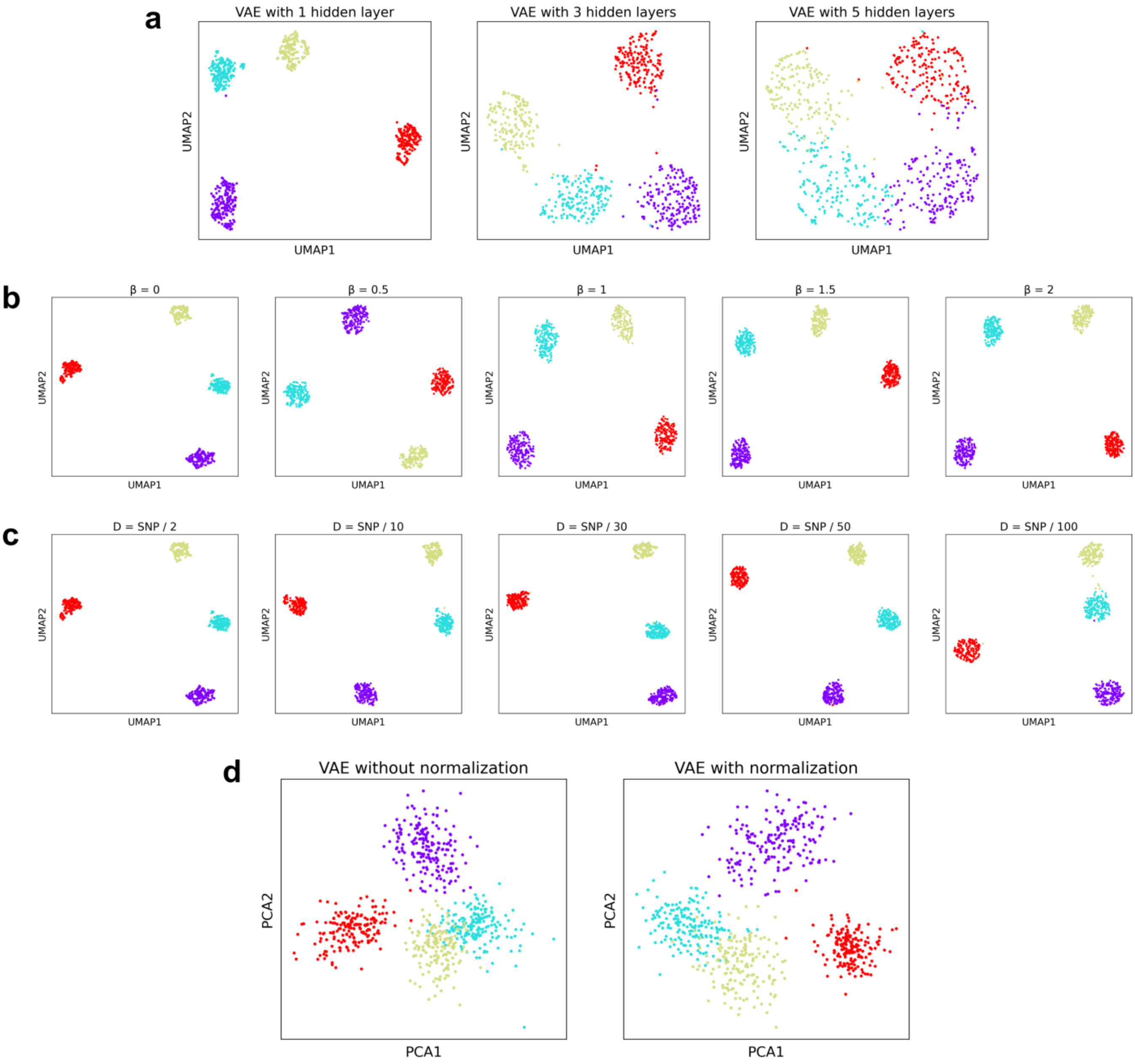


**a,** Genomic geometrical manifolds of Donor4 dataset in UMAPs of the latent space in VAEs with 1, 3, 5 hidden layers. VAE with 1 hidden layer geometrically separates 4 donors the best. **b,** Genomic geometrical manifolds of Donor4 dataset in UMAPs of the latent space in VAEs with 1 hidden layer, and $\beta = 0, 0.5, 1, 1.5, 2$. VAE with $\beta = 0$ geometrically separates 4 donors the best. **c,** Genomic geometrical manifolds of Donor4 dataset in UMAPs of the latent space in VAEs with 1 hidden layer, and D = number of input SNPs divided by 2, 10, 30, 50, 100. VAE with D = 1/2 of the number of input SNPs geometrically separates 4 donors the best. **d,** Principal components of the latent space in VAEs with 1 hidden layer with or without observed-SNP normalization (cell-specific division by the number of observed SNPs: sum(DP > 0)).

Supplementary 3

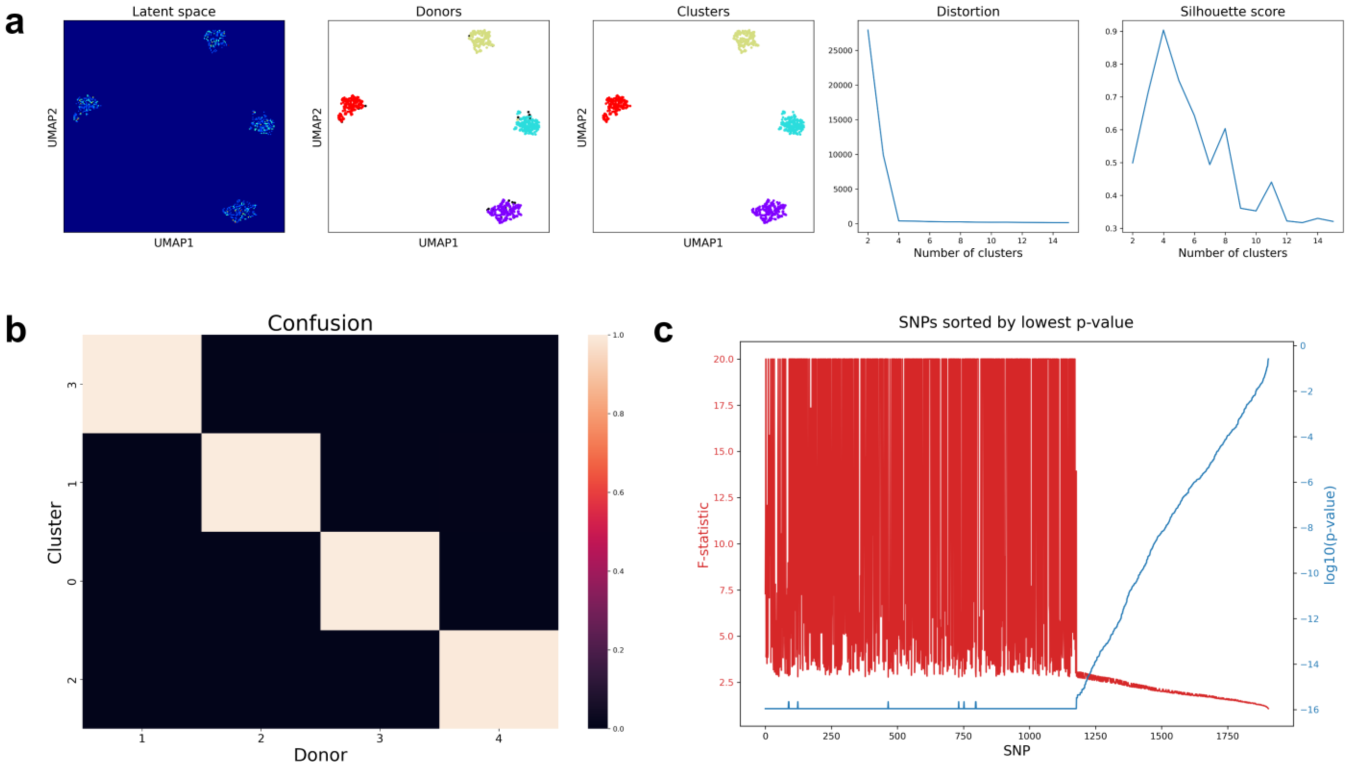


**a,** Density plot of genomic manifold of Donor4 dataset, donor labels and cluster labels on the manifold, distortion and silhouette scores of k-means clustering. There are 4 disconnected manifolds in the density plot corresponding to 4 donors, and k-means clustering identifies them successfully with good clustering metrics. **b,** Confusion matrix between donor labels and cluster labels of Donor4 dataset. The accuracy reaches 99.8%. **c,** F-statistics and p-values of the SNPs of Donor4 dataset ranked by SNPmanifold.

Supplementary 4

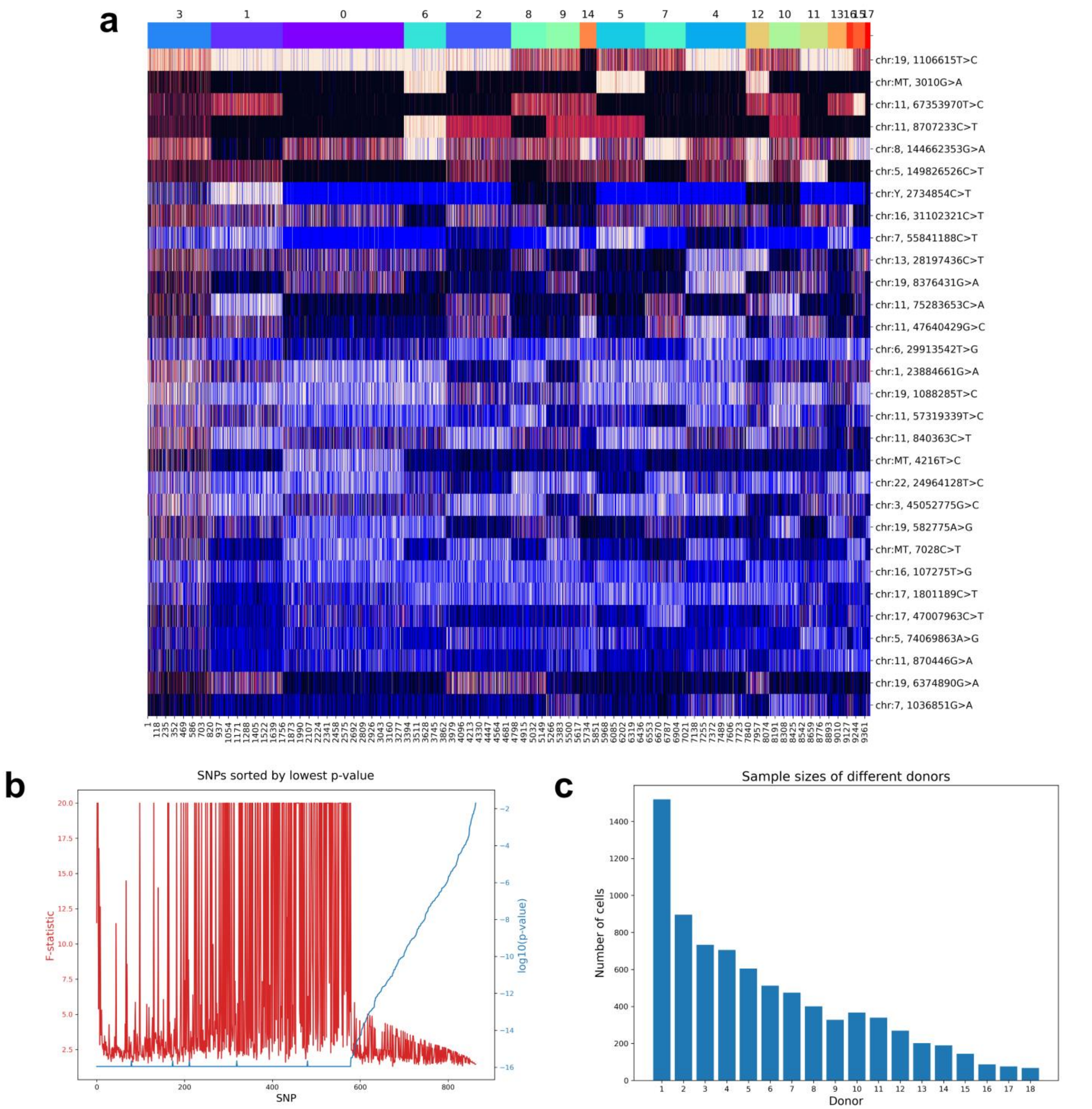


**a,** Heatmap of allele frequency matrix of Donor18 dataset. Blue color indicates missing values (i.e. DP = 0). **b,** F-statistics and p-values of the SNPs of Donor18 dataset ranked by SNPmanifold. **c,** Sample sizes of different donors in Donor18 dataset. The sample size of one donor ranges from 67 to 1519 cells.

Supplementary 5

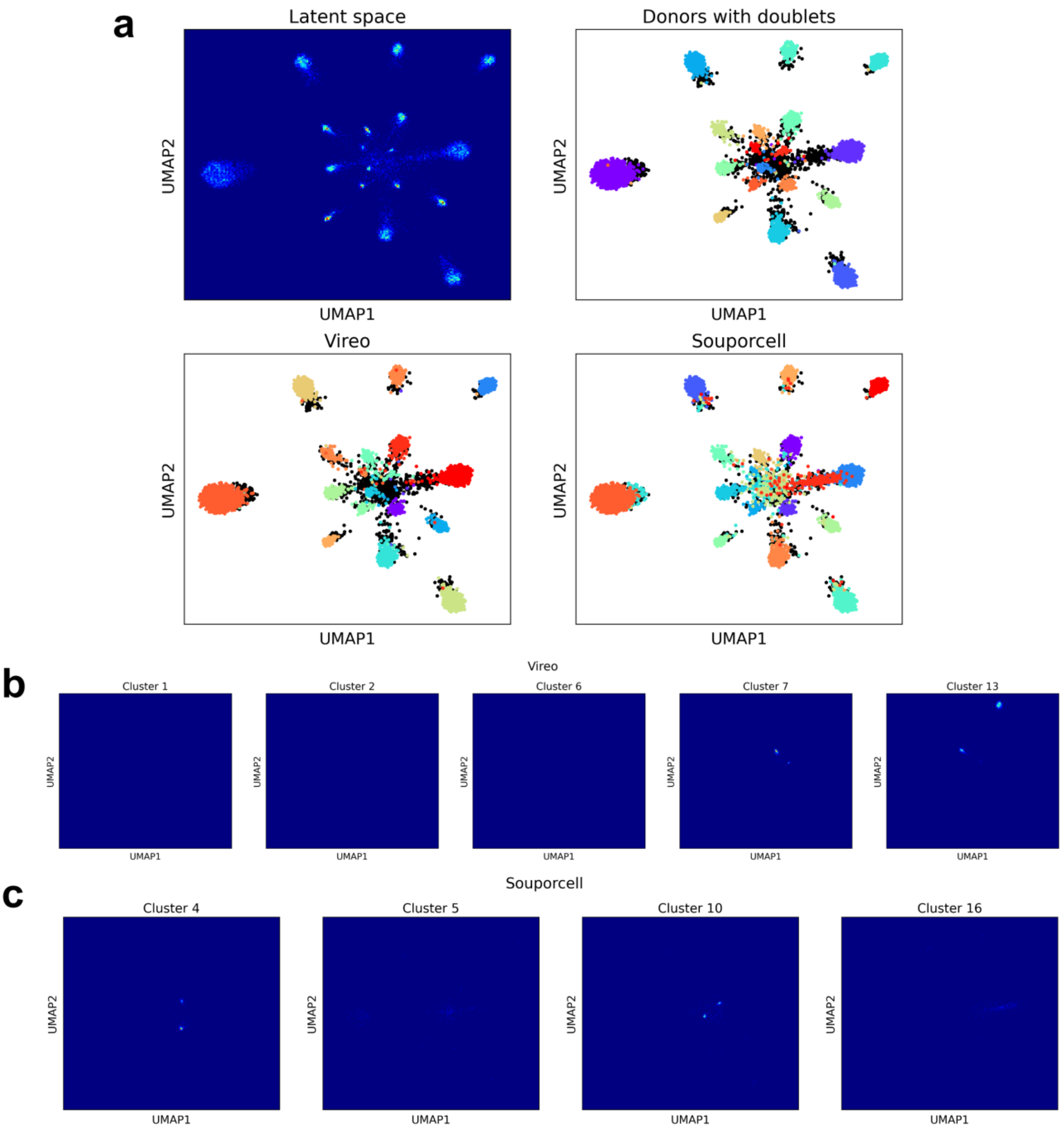


**a,** Density plot of genomic manifold of Donor18 dataset, donor labels with doublets, Vireo labels and Souporcell labels on the manifold. **b,** Abnormal clusters assigned by Vireo without reference to genotypes. Almost no cells are assigned to clusters 1, 2, 6 while 2 donors are combined together in clusters 7, 13. **c,** Abnormal clusters assigned by Souporcell. Parts of doublets are assigned to clusters 5, 16 while 2 donors are combined together in clusters 4, 10.

Supplementary 6

a

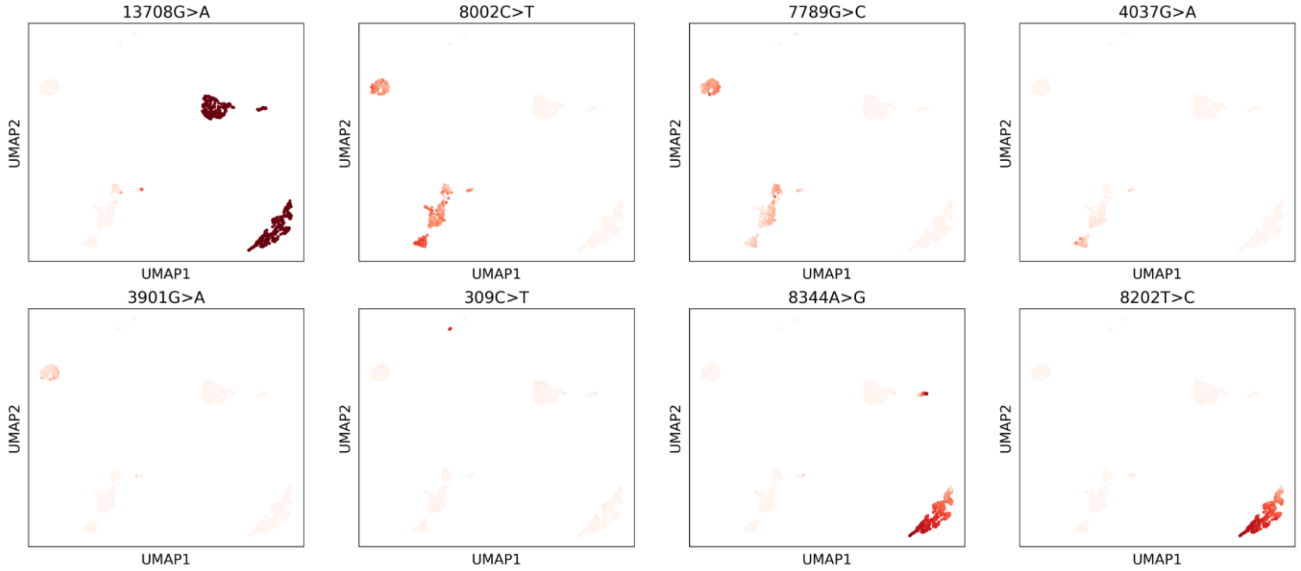


b

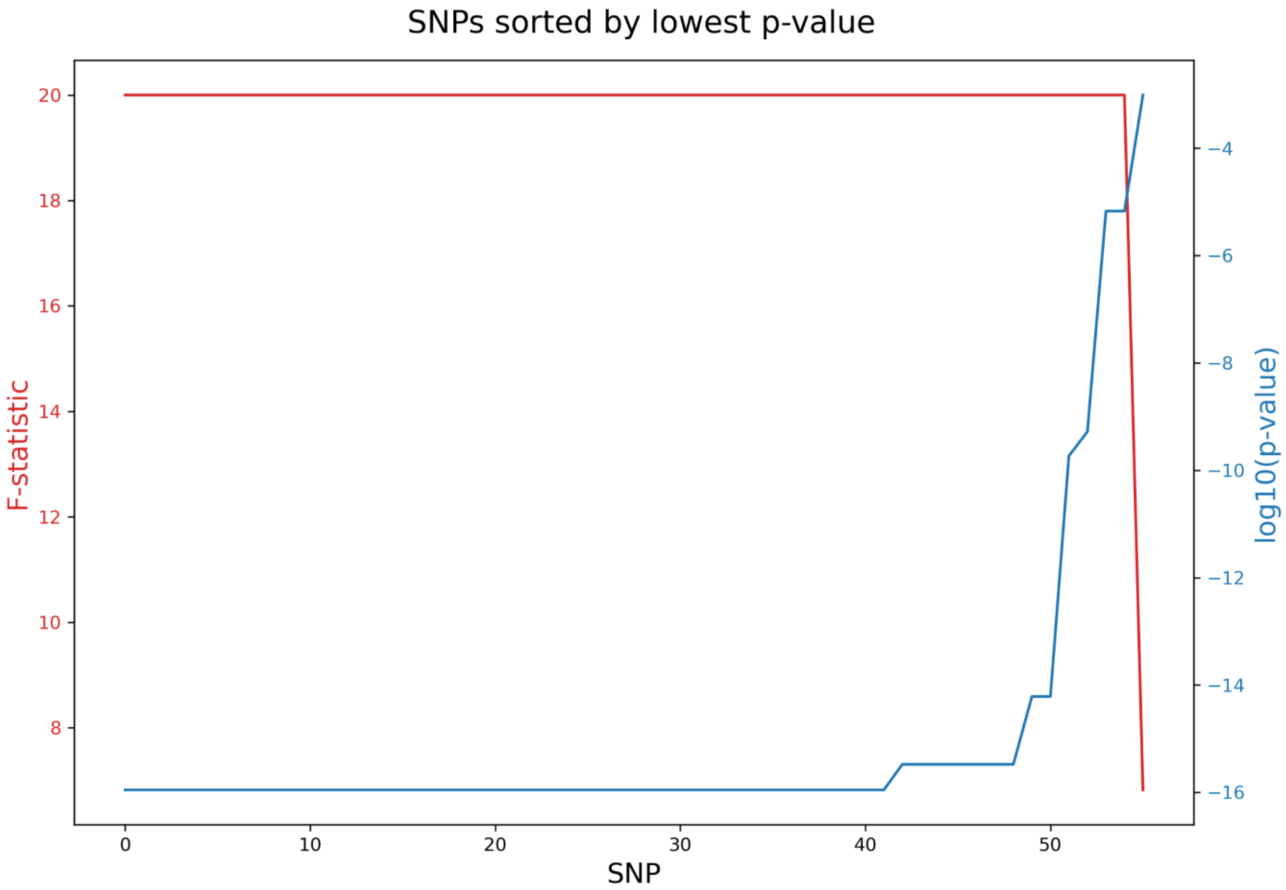


**a,** Allele frequency of 8 SNPs on genomic manifold of TF1_GM11906 dataset. **b,** F-statistics and p-values of the SNPs of TF1_GM11906 dataset ranked by SNPmanifold.

Supplementary 7

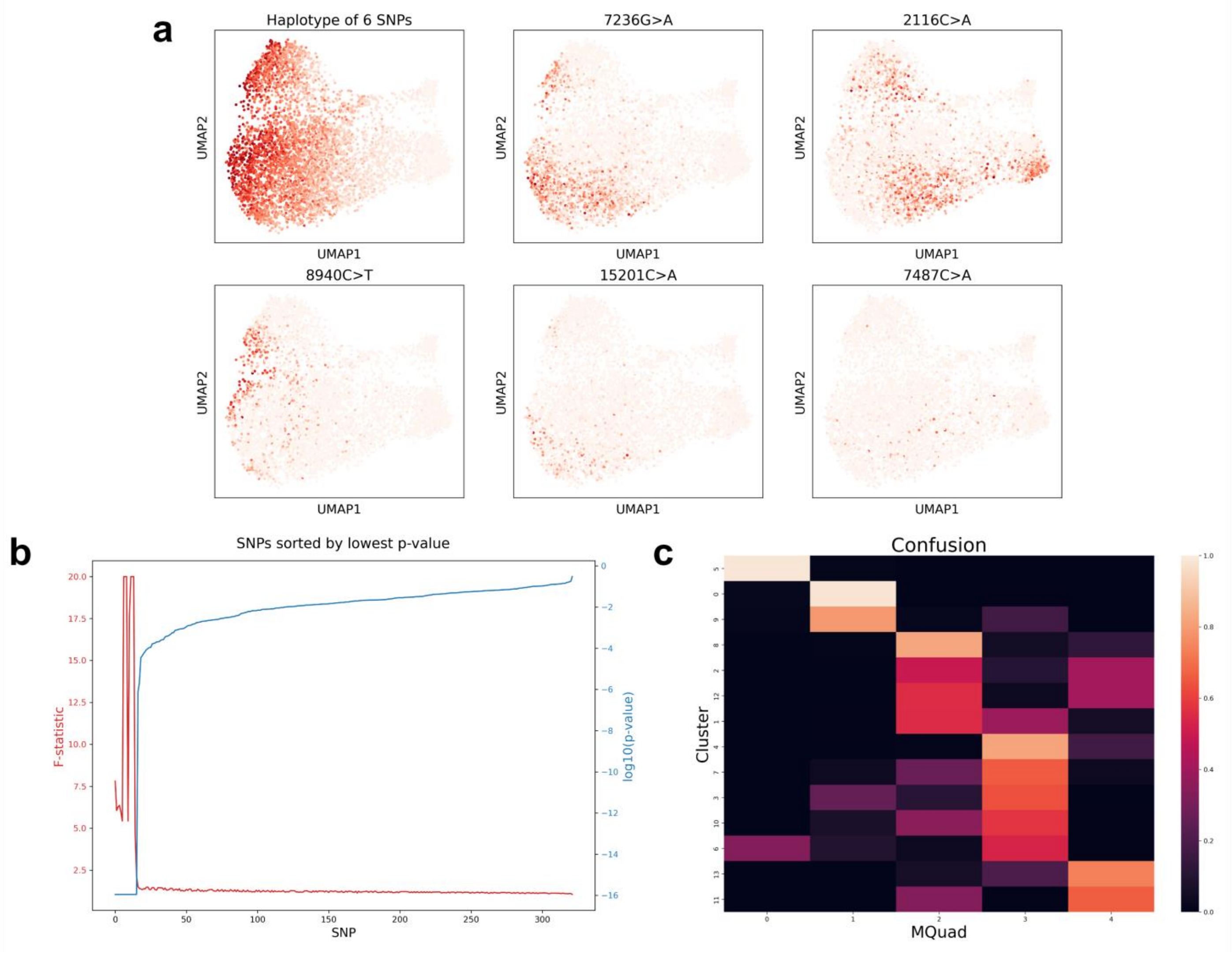


**a,** Allele frequency of 6 mutations on genomic manifold of MKN45 dataset. **b,** F-statistics and p-values of the SNPs of MKN45 dataset ranked by SNPmanifold. **c,** Confusion matrix between MQuad labels and cluster labels of MKN45 dataset. MQuad labels and cluster labels are highly correlated. This means cluster labels of genomic manifold separate MQuad labels into sub-clones with more details.

Supplementary 8

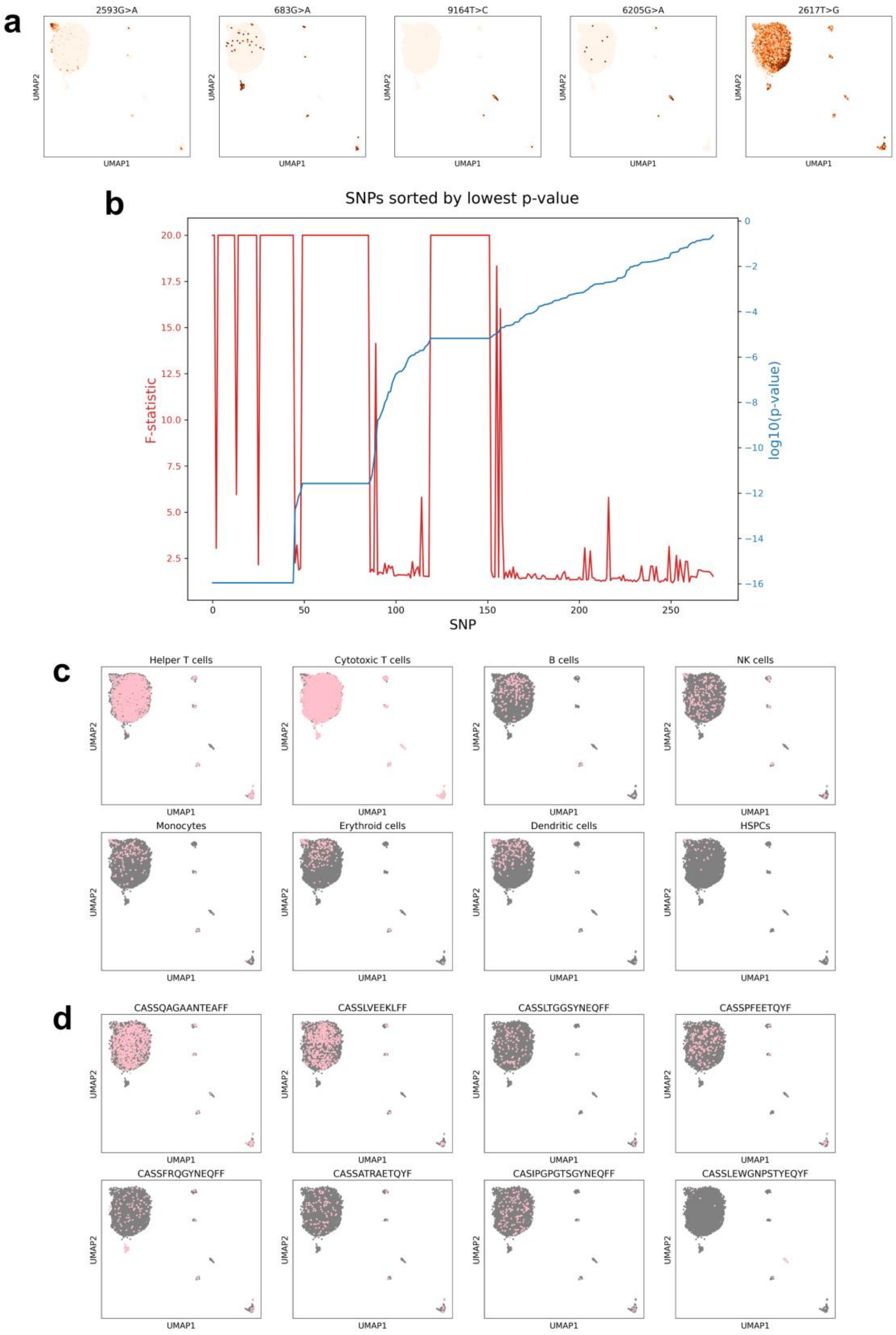


**a,** Allele frequency of 5 SNPs on genomic manifold of BPDCN dataset. **b,** F-statistics and p-values of the SNPs of BPDCN dataset ranked by SNPmanifold. **c,** Cell-type labels on genomic manifold of BPDCN dataset. **d,** TRB-clonotype labels on genomic manifold of BPDCN dataset.

Supplementary 9

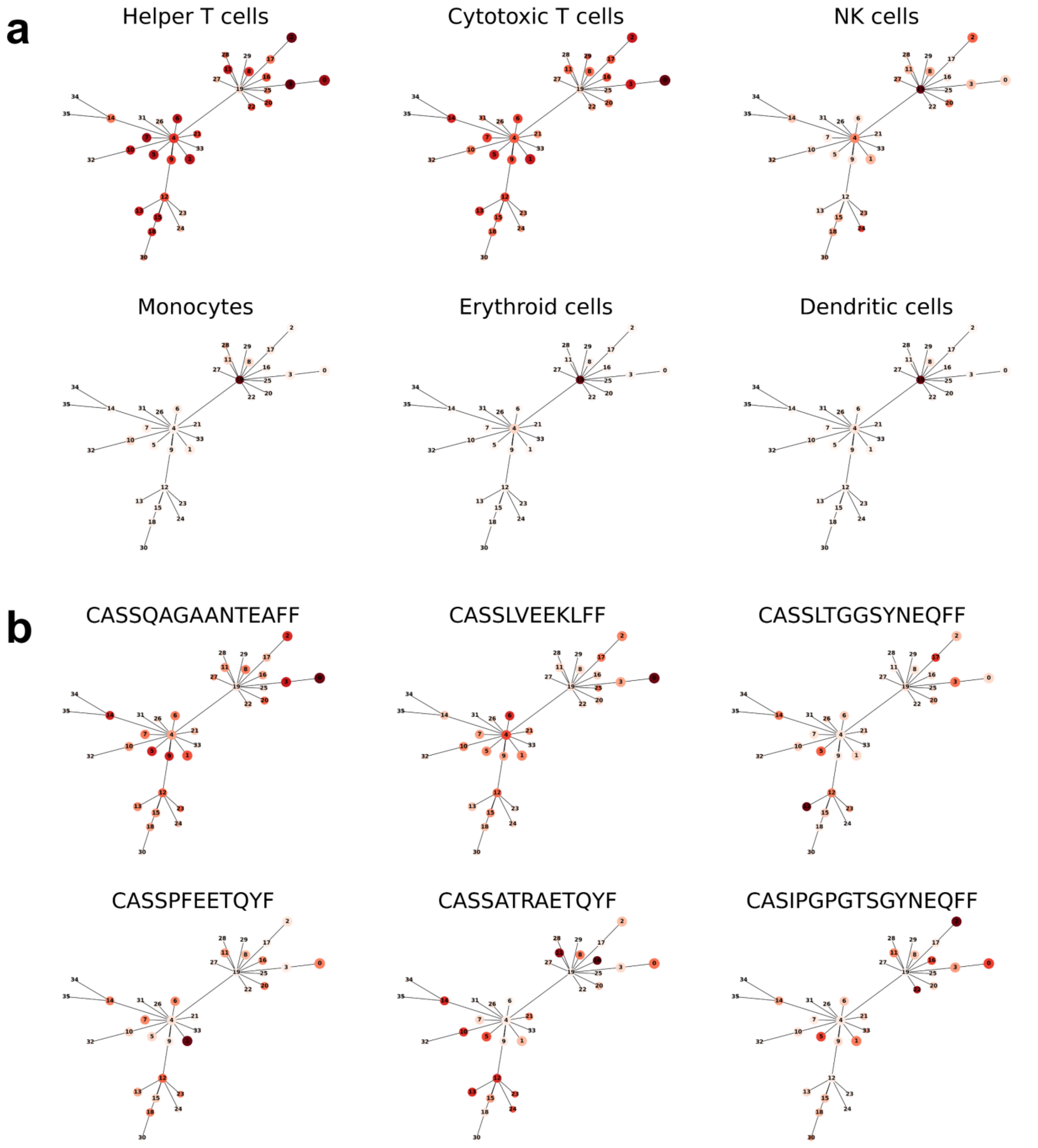


**a,** Relative abundance of 6 cell types on phylogenetic tree of the manifold of BPDCN dataset. Deeper color indicates higher abundance. There are strong correlations between certain cell types and certain clusters. **b,** Relative abundance of 6 TRB clonotypes on phylogenetic tree of the manifold of BPDCN dataset. Deeper color indicates higher abundance. There are strong correlations between certain TRB clonotypes and certain clusters.

Supplementary 10

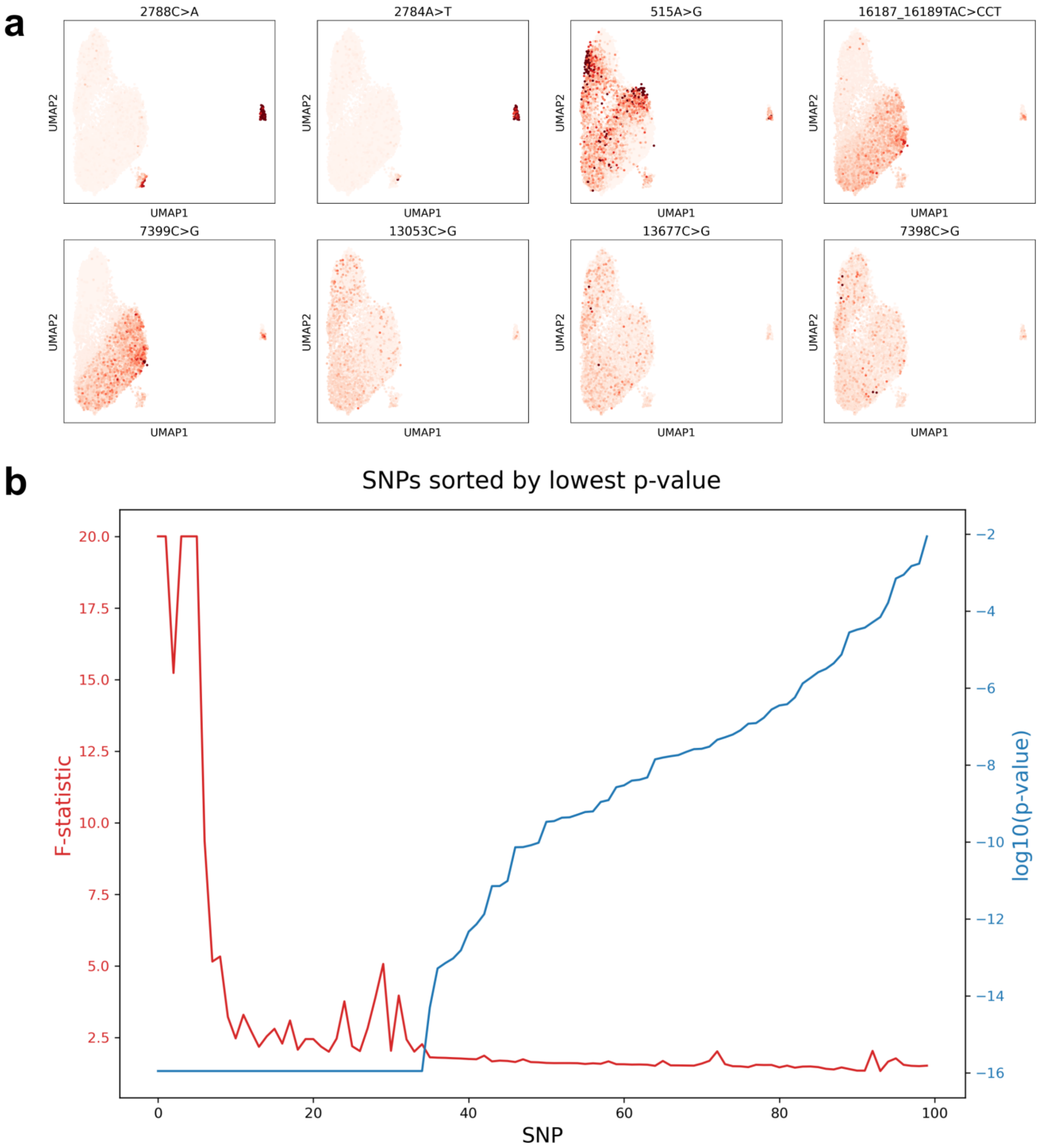


**a,** Allele frequency of 8 mutations on genomic manifold of HSPC_PBMC dataset. **b,** F-statistics and p-values of the SNPs of HSPC_PBMC dataset ranked by SNPmanifold.
